# Vector competence of *Aedes aegypti, Aedes albopictus*, and *Culex quinquefasciatus* mosquitoes for Mayaro virus

**DOI:** 10.1101/661884

**Authors:** Thiago Nunes Pereira, Fabiano Duarte Carvalho, Silvana Faria De Mendonça, Marcele Neves Rocha, Luciano Andrade Moreira

## Abstract

Newly emerging or re-emerging arthropod-borne viruses (arboviruses) are important causes of human morbidity and mortality nearly worldwide. Arboviruses such as Dengue (DENV), Zika (ZIKV), Chikungunya (CHIKV) and West Nile virus (WNV) underwent an extensive geographic expansion in the tropical and sub-tropical regions of the world. In the Americas the main vectors, for DENV, ZIKV and CHIKV, are mosquito species adapted to urban environments namely *Aedes aegy*pti and *Aedes albopictus*, whereas the main vector for WNV is the *Culex quinquefasciatus*. Given the widespread distribution in the Americas and high permissiveness to arbovirus infection, theses mosquito species might pose an important role in the epidemiology of other arboviruses normally associated to sylvatic vectors. Here, we test this hypothesis by determining the vector competence of *Ae. aegypti, Ae. albopictus* and *Cx. quinquefasciatus* to Mayaro (MAYV) virus, a sylvatic arbovirus transmitted mainly by *Haemagogus janthinomys* that have been causing an increasing number of outbreaks in South America namely in Brazil. Using field mosquitoes from Brazil, female mosquitoes were experimentally infected and their competence for dissemination and transmission for MAYV was evaluated. We found high dissemination rate for MAYV in *Ae. aegypti* (57.5%) and *Ae. albopictus* (61.6%), whereas very low rates were obtained for *Cx. quinquefasciatus* (2.5%). Concordantly, we observed that *Ae. aegypti* and *Ae. albopictus* have high transmission ability (69.5% and 71.1% respectively), conversely to *Cx. quinquefasciatus* that is not able to transmit the MAYV. Notably, we found that very low quantities of virus present in the saliva (undetectable by RT-qPCR) were sufficient and virulent enough to guarantee transmission. Although *Ae. aegypti* and *Ae. albopictus* mosquitoes are not the main vectors for MAYV, our studies suggest that these vectors may play a significant role in the transmission of this arbovirus, since both species showed high vector competence in laboratory conditions.

**Author summary:** The present study showed that *Ae. aegypti* and *Ae. albopictus* mosquitoes have high vector competence for MAYV, in laboratory. In contrast, *Cx. quinquefasciatus* mosquitoes were shown to be refractory to MAYV. Regarding the viral dilution and nanoinjection, higher detection sensitivity was observed after virus nanoinjection into naïve mosquitoes, indicating that only a few viral particles are required to infect mosquitoes, and these particles may not be detected by RT-qPCR before the nanoinjection procedure.

## Introduction

The mosquitoes *Ae. aegypti, Ae. albopictus* and *Cx. quinquefasciatus* are widely present throughout the world, especially in tropical and subtropical regions(1–3). They are considered a serious concern to public health as transmission agents of several arboviruses like DENV, ZIKV, CHIKV and WNV (4–13). Studies have shown that *Ae. aegypti* and *Ae. albopictus* mosquitoes exhibit laboratory vector competence for the infection and transmission of MAYV, and *Cx. quinquefasciatus* mosquito has been found infected with MAYV in Cuiabá(14–16).

The MAYV was first isolated in 1954 in rural workers in Mayaro, in the island of Trinidad(17) and as CHIKV, it is an arbovirus of the genus *Alphavirus*, belonging to the family *Togaviridae*(18–20). This arbovirus is transmitted primarily by the bite of female mosquitoes of the genus *Hemagogus*, and in nature several vertebrate can host this virus, being detected in non-human primates, rodents, birds, sloths, and other small mammals(21).

In humans, the MAYV infection usually occurs in people with records of activities in forest areas(18–20,22). The last years in Brazil, several cases of MAYV have been registered in Pará 2008, Mato Grosso 2012 and Goiás 2014–2016(20,22–28). Although there is still no evidence of the transmission efficiency of MAYV in an urban cycle, it has the potential to establish an epidemic scenario in the Americas, similar to what happened to ZIKV and CHIKV(29).

This disease has a similar symptomatology to infection for DENV and/or CHIKV, causing an acute and self-limiting febrile illness, which may be accompanied by hemorrhagic phenomena such as petechiae and gingival bleeding. Usually the wrist, ankle, hands and feet joints are significantly affected and symptoms may persist for several months, incapacitating the infected person(26,30,31).

Using mathematical models taking into account outbreaks since 1960 and increasing global temperature, Lorenz *et al*., 2017(32) predicted that MAYV would expand its area of coverage in the coming years. Thus, the importance of MAYV as a human pathogen with potential to emerge in urban areas is strong. Therefore, the main objective of this study was to evaluate the vector competence of *Ae. aegypti, Ae. albopictus* and *Cx. quinquefasciatus* for MAYV, since these mosquitoes may be involved with the dispersion of this virus.

## Methodology

### Mosquito populations and rearing

For this study, three mosquito species were used: *Ae. aegypti, Ae. albopictus* and *Cx. quinquefasciatus. Ae. aegypti* and *Ae. albopictus* mosquitoes were collected from ovitraps, whereas *Cx. quinquefasciatus* were collected using an entomological ladle. All collections occurred in the neighborhood of Pampulha, in the city of Belo Horizonte, Brazil, in the first half of 2017.

Eggs/larvae were transported to the insectary of René Rachou Institute, Fiocruz/MG, and were kept in a controlled environment, at 27 °C ± 2 °C, ∼82% RH, 12 hr light regime. Larvae were maintained with fish food pellets, Tetramin tropical (Tetra®) and GoldFish colour (Alcon®). After adult emergence and identification, they were kept on 10% sucrose solution regimen, *ad libitum*. Adult females were fed with human blood in an artificial feeder for egg production. To simulate field conditions and minimize the effects of inbreeding all experiments and population colonization, all experiments were conducted with mosquitoes from the same geographic area and up to the fourth generation.

### Mayaro virus culture

An aliquot of MAYV was supplied by the Flavivirus Laboratory of the Oswaldo Cruz Instituto (IOC / Fiocruz); this virus was passaged in *Ae. albopictus* cell line (C6/36), in Leibowitz L-15 medium supplemented with 10% fetal calf serum, and maintained at 28 °C. To prepare the MAYV for oral feeding, a frozen virus stock was passaged once again through C6/36 cells (approximately 2 million cells) and the supernatant was harvested after 5 days (for the first experiment) and at day 4 (for the second experiment).

### Mosquito infection

For the infection, five day-old adult females were allowed to ingest a mixture of viral supernatant and human blood (2:1) which was offered for 45 minutes through glass feeders using pig intestine (from local butcher shop; used as sausage skin) as the membrane and a water jacket system with the temperature maintained at 38 °C. Immediately after feeding, fully engorged females were separated and maintained with a 10% sucrose solution until the end of the experiment.

Two replicates (A and B) were used to evaluate the vector competence of the mosquitoes and in both, fresh viral supernatant was used, from C6/36 cell culture. For the first replicate, the viral titer was 1×10^9^ PFU/mL, and for the second was 6×10^9^ PFU/mL. Both viral titers were quantified after a freezing cycle. Mosquitoes were collected from both groups on different dpi (days post-infection) and stored at −80 °C before processing.

The human blood used in all experiments was obtained from a blood bank (Fundação Hemominas), according to the terms of an agreement with the Instituto René Rachou, Fiocruz/MG (OF.GPO/CCO agreement – Nr 224/16).

### Saliva collection and nanoinjection

At 14 dpi, mosquitoes were submitted to the salivation process. Mosquitoes were anesthetized with CO_2_ and kept on an ice plate while the legs and wings were removed. Each mosquito proboscis was inserted in a 10μL pipette tip containing a 1:1 solution of 10μL of 30% sucrose and sterile fetal calf serum. After 30 minutes, the contents of the tips were individually collected in 0.6 mL tubes and stored at −80 °C until processing.

For the nanoinjection, samples of undiluted saliva from each mosquito group were nanoinjected into 10 to 15 individual naïve *Ae. aegypti*, using a portable injector Nanoject II (Drummond Sci). In each mosquito, a 276nL dose of saliva was nanoinjected intrathoracically (pleural membrane) with a pulled glass capillary. Nanoinjected mosquitoes were collected at 5 days post nanoinjection. The presence of the virus was determined by RT-qPCR analysis; in which on average 6 per group were processed.

### Virus serial dilution and nanoinjection

To better understand our findings around negative heads producing positive mosquitoes after nanoinjection, we tested the sensitivity of virus detection particles through RT-qPCR. Serial dilutions of known virus stocks were nanoinjected into naïve mosquitoes. Replicates A and B had an initial concentration of 2.09×10^4^ and 5.59×10^4^, respectively, and were further diluted 6 times. As control, uninfected cell supernatant was used.

### Analysis of MAYV by RT-qPCR

In this study, we used the head+thorax of orally-infected mosquitoes to verify the dissemination rate and whole mosquitoes to verify the transmission/infectivity of saliva. Total RNA from head+thorax, whole body mosquito or viral supernatant was extracted with the High Pure Viral Nucleic Acid Kit (Roche), following the manufacturer’s instructions. The thermocycling conditions were as follows: reverse transcription at 50 °C for 10 min; RT inactivation/initial denaturation at 95 °C for 30s, 40 cycles of 95 °C for 5s and 60 °C for 30s, followed by cooling at 37 °C for 30s. The volume of the reaction was 10μL (5× LightCycler® Multiplex RNA Virus Master (Roche), 1μM primers and 0.2μM probe and 125 ng RNA.

MAYV infection/dissemination in mosquitoes was quantified by RT-qPCR using LightCycler® 96 (Roche). A multiplex assay was performed according to previous studies(14,33,34). All samples were tested in duplicate for MAYV and the viral titer was determined through standard curve using serial dilutions of the target gene cloned into the pGEMT-Easy plasmid (Promega).

## Data analysis

Mosquito infection data were analyzed with the Person omnibus normality and D’Agostino tests. Fisher’s exact test was then used to assess differences in viral prevalence. Comparisons were significant for *P* values lower than 0.05 and viral load data were compared using a Mann-Whitney U test. All analyses were performed using Prism V 7.4 (Graphpad).

## Results

### MAYV dissemination in Ae. aegypti, Ae. albopictus and Cx. quinquefasciatus

Our results showed that at the analyses post-infection times, *Ae. aegypti* and *Ae. albopictus* became positive for MAYV. In addition, in both replicates (A and B) there was increased dissemination at 14 dpi compared to 7 dpi in *Ae. aegypti* and *Ae. albopictus.* On the other hand, the *Cx. quinquefasciatus* mosquitoes were shown to be refractory to MAYV.

Between replicates A and B (Fig. 1), out of 40 *Ae. aegypti* mosquitoes fed with a mixture of blood and viral supernatant, viral dissemination at 7 dpi occurred in 17 mosquitoes (42.5%), whereas 57.5% were uninfected. At 14 dpi, viral dissemination occurred in 72.5% of the specimens, while 27.5% were not negative.

**Figure 1.**
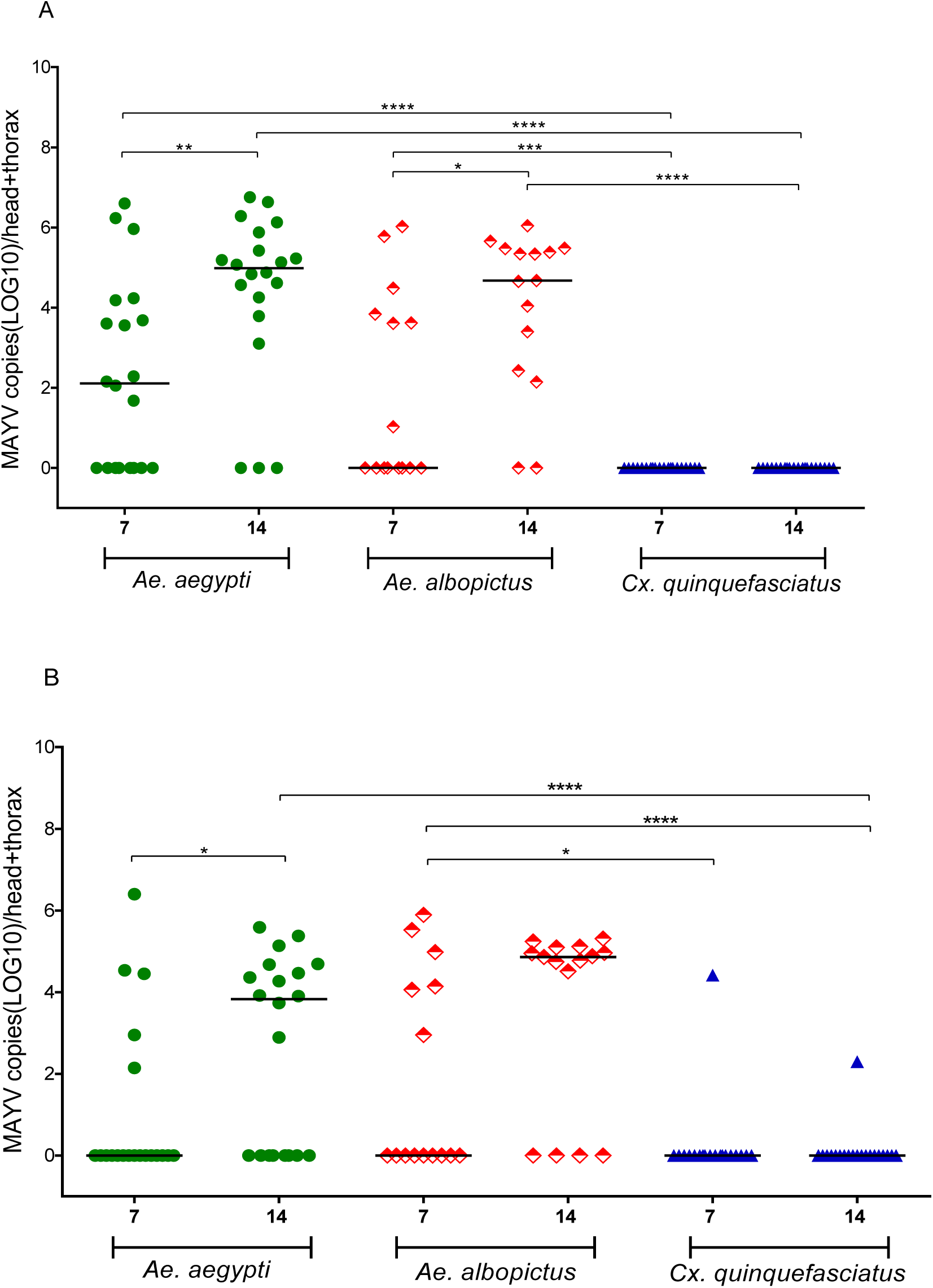
Mosquito viral dissemination. (replicates A and B) – Each point represents a single adult female and the black lines indicate the median of the Mayaro virus copies in each group. The asterisks represent P <0.005 after the Mann-Whitney U-Test.

For *Ae. albopictus* of replicates A and B (Fig. 1), out of 30 mosquitoes evaluated at 7 dpi, viral dissemination was present in 13 (43.3%), whereas 17 (56.66%) were not infected. At 14 dpi, out of 30 *Ae. albopictus*, 24 (80%) were positive and 6 (20%) were uninfected.

The vector competence was also analyzed in *Cx. quinquefasciatus* mosquitoes and there was only mosquito dissemination for MAYV in replicate B (Fig. 1). Out of 40 *Cx. quinquefasciatus* mosquitoes at 7 dpi, the dissemination was present in only 1 sample (2.5%), whereas 39 (97.5%) were not infected. At 14 dpi, out of 40 *Cx. quinquefasciatus*, the dissemination was detected in 1 (2.5%), and 39 (97.5%) were uninfected.

In general, our results show a higher susceptibility for MAYV in *Ae. albopictus* (61.6%), followed by *Ae. aegypti* (57.5%). However, *Cx. quinquefasciatus* mosquitoes may be considered refractory to MAYV infection (only 2.5%).

Pairwise comparisons showed that for replicate A, there were differences between the groups at 7dpi: *Ae. aegypti* × *Cx. quinquefasciatus* (*P* <0.0001), *Cx. quinquefasciatus* × *Ae. albopictus* (*P* = 0.0010); and at 14 dpi: *Ae. aegypti* × *Cx. quinquefasciatus* (*P* <0.0001), *Cx. quinquefasciatus* × *Ae. albopictus* (*P* <0.0001), between 7 × 14 dpi for *Ae. aegypti* (*P* = 0.00043) and 7 × 14 dpi for *Ae. albopictus (P* = 0.0230). For replicate B, there were differences between the groups at 7dpi for *Cx. quinquefasciatus* × *Ae. albopictus* (*P* = 0.0207) and at 14 dpi for *Ae. aegypti* × *Cx. quinquefasciatus* (*P* <0.0001), *Cx. quinquefasciatus* × *Ae. albopictus* (*P* <0.0001) and between 7 × 14 dpi for *Ae. aegypti* (*P* = 0.0262). The dissemination rate for by MAYV, for each species in both experiments and at different dpi, is shown in Table 1.

**Table 1.** *Aedes aegypti, Aedes albopictus* and *Culex quinquefasciatus* orally infected with Mayaro virus. Initial viral titer was determined by plaque formation unit. Infected/total mosquito numbers are shown parenthesis.

| MAYV | Titer<br>(PFU/ml) | Days post<br>infection | *Ae.*<br>*aegypti* | *Ae.*<br>*albopictus* | *Cx.*<br>*quinquefasciatus* |
| --- | --- | --- | --- | --- | --- |
|  |  |  | Head/thorax dissemination rate % |  |  |
| Replicate A | 1×10<sup>9</sup> | 7 | 60 (12/20) | 46.6 (7/15) | 0 (0/20) |
|  |  | 14 | 85 (17/20) | 86.6 (13/15) | 0 (0/20) |
| Replicate B | 6×10<sup>9</sup> | 7 | 25 (5/20) | 40 (6/15) | 5 (1/20) |
|  |  | 14 | 60 (12/20) | 73.3 (11/15) | 5 (1/20) |

### Transmission of MAYV through saliva injection into naïve mosquitoes

To verify if infected mosquitoes were able to transmit MAYV, saliva from positive head+thorax was nanoinjected into naïve Br mosquitoes. Out of 69 *Ae. aegypti* nanoinjected mosquitoes, 48 (69.5%) became infected with MAYV. For *Ae. albopictus* after injection of 59 mosquitoes, 42 (71.1%) became infected with MAYV (Fig 2B). These results show that both mosquitoes have high dissemination rates and transmission capacities for MAYV. It is important to say that more than 80% of all saliva samples (from *Ae. aegypti* and *Ae. albopictus*), were infective to naïve mosquitoes.

**Figure 2.**
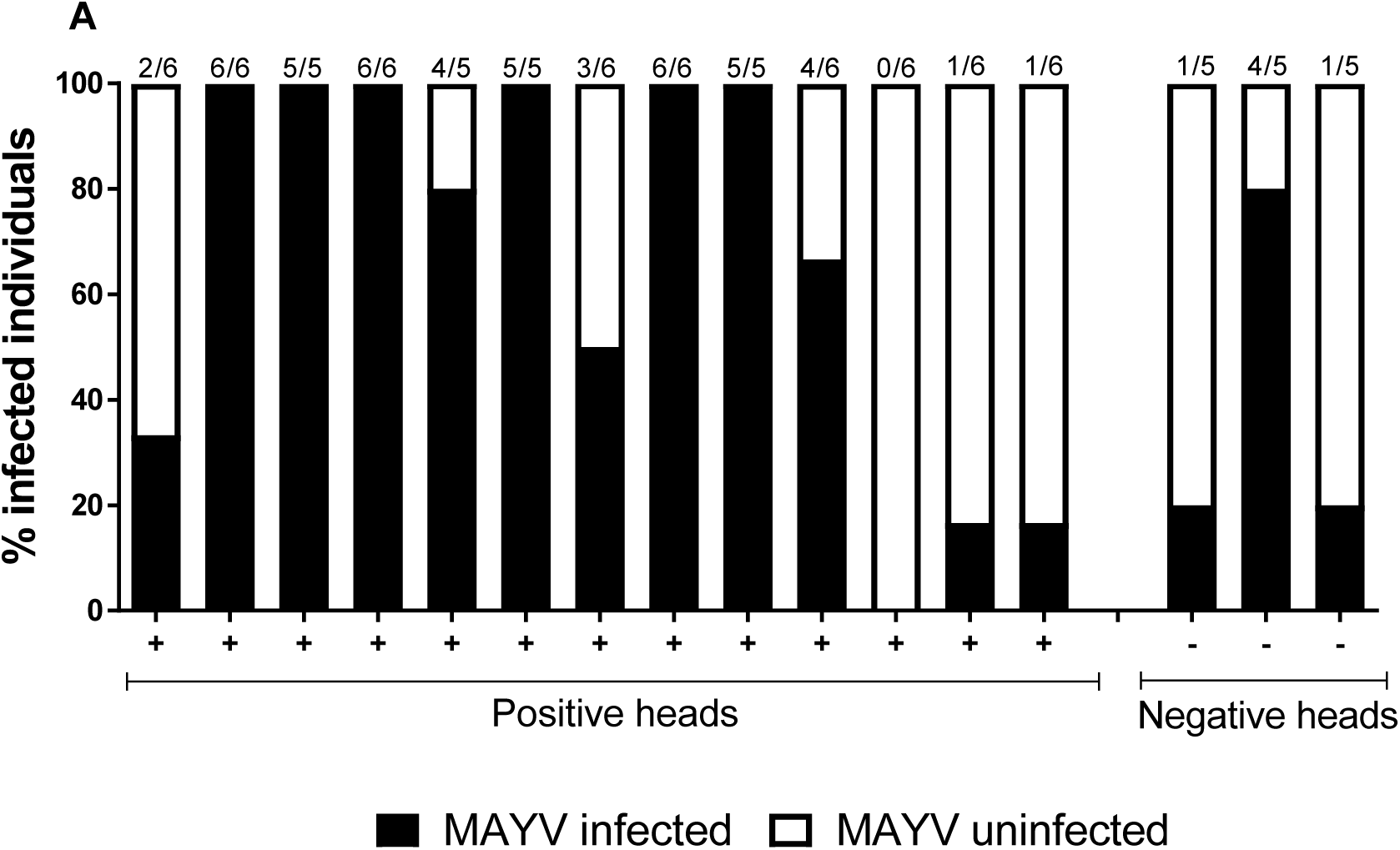
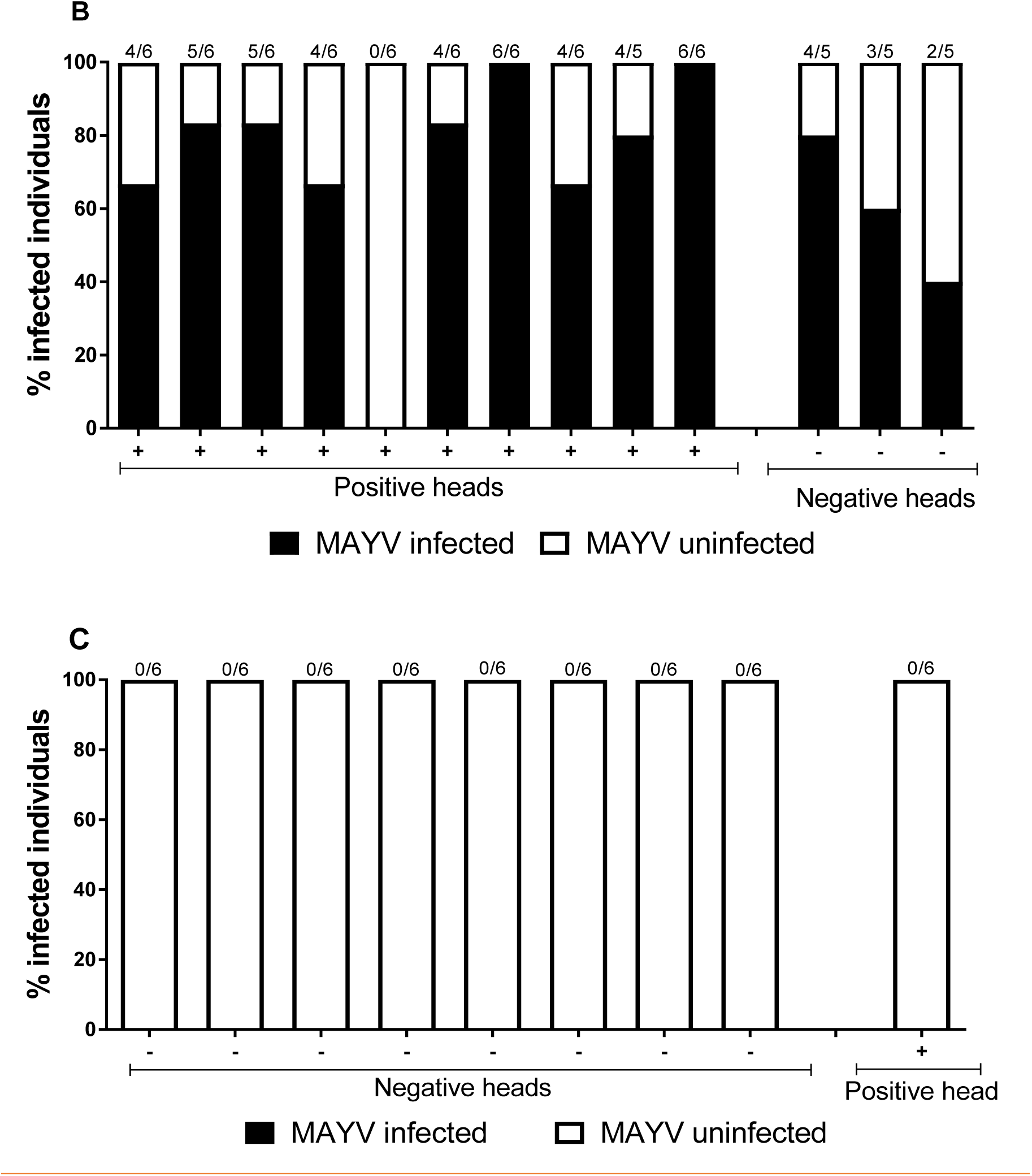
Nanoinjection of saliva from three infected mosquito species into naïve *Aedes aegypti* mosquitoes. Saliva samples were collected from *Ae. aegypti* (A), *Ae. Albopictus* (B) and *Cx. quinquefasciatus* (C) which were previously infected with MAYV, followed by injection into naïve mosquitoes. Infected mosquitoes are shown in black and uninfected in white. Each bar represents a single saliva sample and the number of infected mosquitoes /total nanoinjected mosquitoes is given at the top of each bar.

In contrast, none of the 54 mosquitoes nanoinjected with *Cx. quinquefasciatus* saliva were able to become infected. This was expected as the rate of MAYV infected mosquitoes were very low, based on virus dissemination in head+thorax.

We also nanoinjected saliva originated from negative head+thorax negative *Ae. aegypti* and *Ae. albopictus* mosquitoes and, surprisingly, some samples were able to infect naïve mosquitoes. Saliva samples from 3 negative *Ae. aegypti* head+thorax (Fig.2A) were nanoinjected into 15 mosquitoes, and 6 (40%) became infected with MAYV. Saliva from 3 negative *Ae. albopictus* head+thorax (Fig.2B) was nanoinjected in 15 mosquitoes, and 9 (60%) became infected with MAYV.

### Virus serial dilution and nanoinjection

In order to try to understand why negative heads+thoraces are able to produce infectious saliva we performed a series of virus dilution and injection into naïve mosquitoes, followed by RT-qPCR detection. We found that even when diluted viral samples were not detected through RT-qPCR, they were able to produce positive mosquitoes when nanoinjected (Fig. 3, samples 5A and 6A replicate A and sample 5A replicate B).

**Figure 3.**
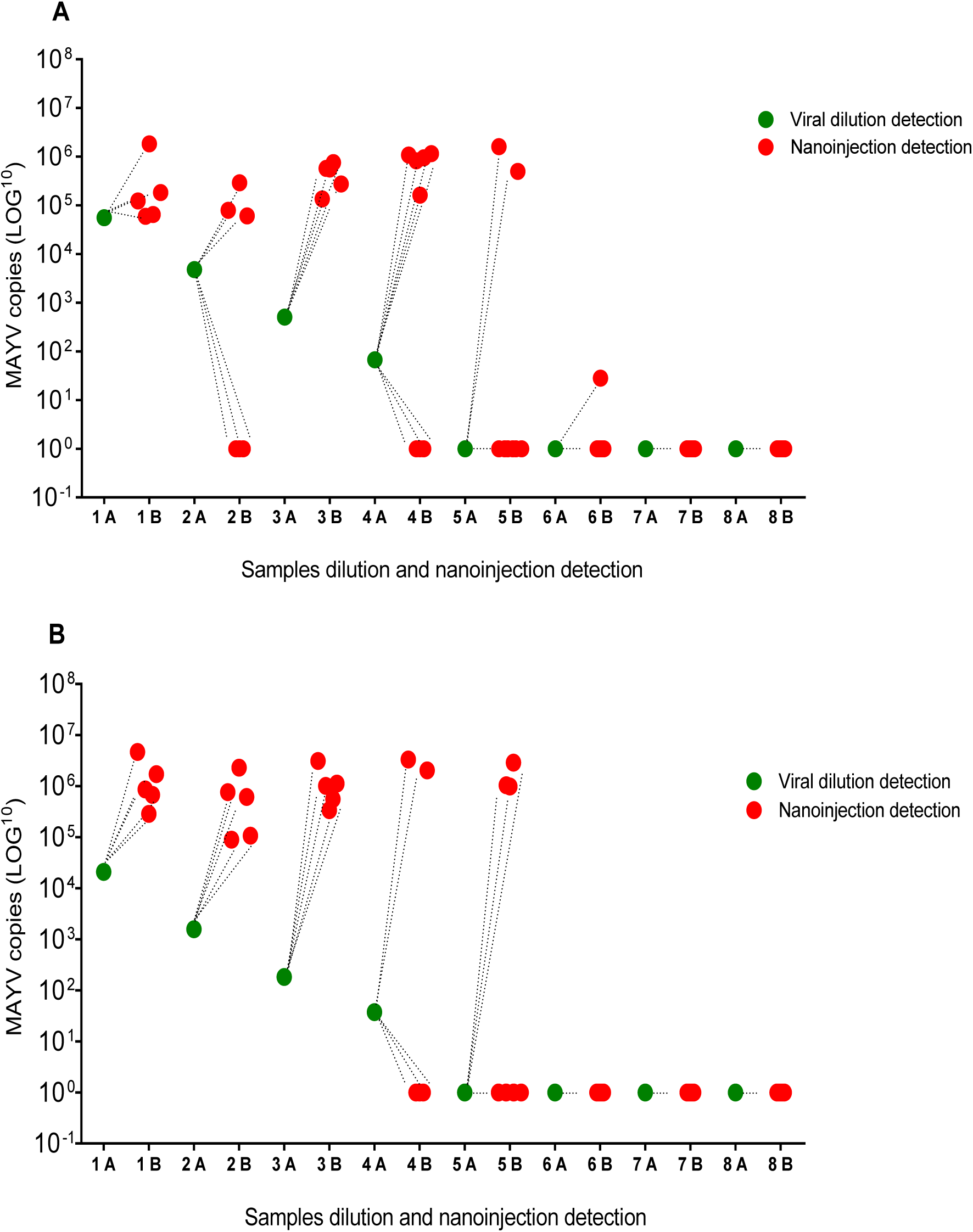
Sample dilutions and detection of Mayaro virus. A and B represent different replicates. Each viral dilution (represented by green dots) was nanoinjected into naïve mosquitoes, followed by virus detection through RT-qPCR (red dots). Samples 8A and 8B are mock control. Each red spot represents a single female mosquito.

The nanoinjection of the viral saliva/dilution in the mosquito acts as a model that amplifies the particles and facilitates detection(35). This may explain why saliva from heads classified as negative produce positive mosquitoes, upon infection. We believe that only a few viral particles are required to infect a mosquito, which cannot be detected through RT-qPCR.

## Discussion

To date, this is the first study to examine the vector competence of *Ae. aegypti, Ae. albopictus* and *Cx quinquefasciatus* mosquitoes for MAYV. Our results show that *Ae. aegypti* mosquitoes are easily infected with MAYV. In general, these mosquitoes presented high levels of infection in heads+thoraces at 14 dpi, averaging 9.6×10^4^ and 4.7×10^4^ viral particles, for replicates A and B, respectively. A previous study by Pereira *et al*., 2018(34), showed that *Ae. aegypti* mosquitoes from Rio de Janeiro were highly susceptible to MAYV, presenting a higher number of viral copies than those obtained in this study. However, this difference may be related to the viral input at the time of infection or even the genetics of the mosquitoes used in our study. The influence of viral dosage is already well described in the virus-vector interaction, and it has been previously reported that *Ae. aegypti* mosquitoes infected with high doses of MAYV present higher infection rates(14,36). As for the genetic variability, Gokhale *et al*., 2015(37) suggested that for CHIKV, which is a similar virus to MAYV(38), the genetic background of the vector strongly influenced the susceptibility to the virus.

MAYV cases have been reported in Brazil, in the North, Northeast and Center-West regions(22–27). *Ae. aegypti* mosquitoes naturally infected with MAYV have been found in Cuiabá, Mato Grosso (15), but little is known about its role of transmission in the urban cycle. Thus, to evaluate viral transmission, MAYV infected saliva of *Ae. aegypti* was nanoinjected into naïve mosquitoes, with 69% of mosquitoes becoming infected. Other laboratory studies also confirm the ability of this vector to transmit MAYV(14,16,34). Therefore, our results confirm that this vector has a great potential for infection and transmission of MAYV and therefore, play an important role in the transmission of this virus, if it gets urbanized. In relation to the *Ae. aegypti* mosquito, Pereira *et al*., 2018(34), observed that when these mosquitoes harbor *Wolbachia* they had significant impaired ability to infect/transmit MAYV. This strategy could become an effective tool for reducing MAYV transmission, throughout Latin America.

The *Ae. albopictus* mosquitoes were found to have a similar infection rate to *Ae. aegypti*, showing high susceptibility to MAYV. At 14 dpi, significant numbers of viral particles were observed in this species. So far, there are few studies showing the relationship between MAYV and *Ae. albopictus*.

Smith and Francy, 1991(39), evaluated the vector efficiency of a Brazilian *Ae. albopictus* mosquito line fed on viremic hamster blood for MAYV and found that the infection ranged from 9% to 16%. The authors have classified this strain as being relatively refractory to MAYV infection, but have suggested that it may become more susceptible, serving as a secondary vector in an outbreak or as a bridge vector between MAYV transmission cycles. Wiggins *et al*., 2018(16), observed in an oral infection experiment that *Ae. albopictus* mosquitoes had a significantly higher infection rate when compared to *Ae. aegypti* mosquitoes, which is similar to those observed in our study.

Oral infection was significantly higher in Ae. albopictus (85–100%) than in *Ae. aegypti* (67–82%). Oral infection was significantly higher in *Ae. albopictus* (85–100%) than in *Ae. aegypti* (67–82%). The same mosquito species may present differentiated vector competence when at different sites, since different genotype/genotype interactions between virus and vector may occur. As an example, it has been demonstrated that the reemergence of the CHIKV may have been facilitated by the genetic adaptation of the virus to *Ae. albopictus* vector(40,41).

In 2016, the first imported case of MAYV in a French citizen was reported in an area where the *Ae. albopictus* mosquito is well established(42), thus highlighting the need to better understand the vector competence of this mosquito, as well as its possible role in the transmission of MAYV.

To evaluate viral transmission in our experiments, *Ae. albopictus* mosquito saliva submitted to MAYV infection were nanoinjected into naïve mosquitoes, resulting in a high rate of infectivity. Smith and Francy, 1991(39), noted in their study that approximately half (5/11) of the mosquitoes infected with hamster viremic blood were able to transmit MAYV when their saliva was tested in capillary tubes. Wiggins *et al.*, 2018(16) observed that *Ae. albopictus* mosquitoes exhibited low rates of MAYV infection in saliva expectorates. However, these authors used a different methodology.

We also attempted to study whether saliva originating from negative head+thorax samples were able to infect naïve mosquitoes. Unexpectedly, negative head+thorax samples from *Ae. aegypti* and *Ae. albopictus* mosquitoes were able to infect other mosquitoes through their saliva. Previous experiments in our group have shown the same effect for DENV with negative thorax and positive saliva, but it resulted in a lower infection rate (date not shown).

To better understand these findings, we investigated the relationship between the amounts of (detectable) particles required for mosquito infection. It was possible to observe that samples classified as negative for RT-qPCR were able to infect other mosquitoes, indicating that only a few viral particles are necessary to initiate infection. In addition, it was observed that regardless of nanoinjected viral doses (10^4^, 10^3^, 10^2^ and 10^1^); they produced an average viral infection between 10^5^ and 10^6^. We suggest that for a broader understanding of this finding, complementary studies using immunofluorescence assays to detect transmission, through saliva, originating from PCR negative heads, should be undertaken.

When evaluating the vector competence of *Cx. quinquefasciatus*, only two mosquitoes were found positive for MAYV, and no mosquito became infected after injection with saliva from these *Cx. quinquefasciatus* mosquitoes submitted to MAYV infection. Corroborating with our data, Brustolin *et al*., 2018(43), also tested the vector competence for *Cx. quinquefasciatus* and observed that this mosquito had either poor or null infection and transmission rates for MAYV.

*Cx. quinquefasciatus* is quite abundant in Brazil and, to date there is only a single record of this mosquito harboring MAYV in Cuiabá(15). This mosquito is, however, a vector of *Wuchereria bancrofti* in Brazil, an etiologic agent of lymphatic filariasis in humans(44), and was recently incriminated in ZIKV transmission in the metropolitan region of Pernambuco (8). Guo *et al*., 2016(45), have also demonstrated the vector competence of *Cx. quinquefasciatus* for ZIKV in China. In contrast, several other studies have demonstrated the lack of ability of this mosquito to infect and transmit ZIKV (46–48). These results confirm that the vector competence of the same mosquito species can vary geographically, emphasizing the importance of studying the vector competence of different mosquitoes and from different localities.

In conclusion, our studies show that *Ae. aegypti* and *Ae. albopictus* can be infected and, potentially transmit MAYV, although *Cx. quinquefasciatus* exhibited poor vector competence. *Ae. aegypti* and *Ae. albopictus* are widely distributed in the Americas(3,49) and although they are not the main vectors for MAYV, they can potentially play a significant role in the transmission of this virus.

## Acknowledgments

We wish to thank you Dra. Ana Maria Bispo de Filippis (IOC – FIOCRUZ) for donating the MAYV isolate. We are in debt to Dr. Marco Antônio Silva Campos and Dr. Alexandre de Magalhães Vieira Machado and their team, who provided viral culture infrastructure in the Laboratório de Imunologia de Doenças Virais (IRR – FIOCRUZ). We are grateful to all members from the Group Mosquitos Vetores: Endossimbiontes e Interação Patógeno-Vetor, (IRR – FIOCRUZ), especially for Dr. Alvaro Gil Araujo Ferreira for his critical reading of the manuscript. We are grateful to Belo Horizonte municipality and the Universidade Federal de Minas Gerais who helped to collect mosquito samples. This study was financed in part by the Coordenação de Aperfeiçoamento de Pessoal de Nível Superior – Brasil (CAPES) – Finance Code 001”, FAPEMIG, CNPq and, indirectly, by the World Mosquito Program.

## Author contribution

T.N.P., and F.D.C., contributed to the conception and design of all the experiments. T.N.P., S.F.M., M.N.R., and F.D.C., were involved in the development of the experiments. T.N.P., analyzed the data and drafted the manuscript. L.A.M., supervised the research. All authors read and approved the final manuscript.

